# 3D-UHU-TU: A Three-Dimensional Bladder Cancer Model in a Healthy Urothelial Environment

**DOI:** 10.1101/2024.10.22.619472

**Authors:** Benjamin O. Murray, Jinhui Gao, Katherine Swarbrick, Alex Freeman, Jennifer L. Rohn

## Abstract

Bladder cancer cases and fatalities continue to rise worldwide with treatment outcomes not improving in the last four decades. Poor translation of potential new therapies from pre- clinical studies to the clinic could be one reason behind this. The patient-derived xenograft (PDX) mouse is the gold-standard for testing new bladder cancer therapies, but there are key physiological and molecular differences between mouse and human bladders. Thus, more human cell-based models may improve translation of treatments.

Here, we introduce a bladder cancer microtissue model called <u>3D</u> <u>U</u>rine-tolerant <u>Hu</u>man <u>U</u>rothelium-Tumour (3D-UHU-TU), which incorporates spheroids derived from human bladder cancer cell lines RT112 (low grade) and T24 (high grade) into the previously published 3D-UHU healthy urothelial model in a 100% urine environment. Both low- and high-grade 3D- UHU-TU models were characterised using immunofluorescence and immunohistochemistry staining with diagnostic markers (CK7, CK20 and GATA3), cadherin markers (E- and N-Cadherin), invasion and migration markers (MMP-2 and MMP-9) and a proliferation marker (Ki-67). Both models expressed the correct markers in the correct spatial areas. We also investigated the utility of both 3D-UHU-TU models as a platform to test treatments, using the conventional chemotherapeutic Mitomycin C as proof of principle. After 2 hours of treatment and 24 hours of recovery, cell lysis and nuclear damage were observed in both low- and high-grade cancer spheroids, with minimal damage to the surrounding healthy urothelium. At higher doses, cancer spheroids either disintegrated or were reduced in size, with the healthy urothelium still intact.

Taken together, 3D-UHU-TU is a novel, *in vitro* model for testing both the safety and efficacy of new treatments. Furthermore, our work lays the foundation for testing treatments on patient-derived tumour spheroids in a personalised medicine approach.

## 1. Introduction

Bladder cancer is one of the most expensive cancers to treat (1, 2), and cases and deaths continue to rise year on year (3, 4). Due to the heterogenous nature of bladder cancer (5), treatment strategies specific to the particular tumour type are limited. This may explain in part why treatment outcomes have not improved for the last four decades (6), lagging behind progress with many other cancers (7).

There are two main types of urothelial carcinoma: non-muscle invasive (NMIBC) and muscle-invasive bladder cancer (MIBC). NMIBC (clinical stage Tis, Ta and T1) is the most common (∼75% of cases), which can be further separated into low-, intermediate- and high-risk categories, for which different treatments can be implemented depending on tumour grade (8). First-line treatments include transurethral resection of the bladder tumour (TURBT), Bacillus Calmette-Guerin (BCG) immunotherapy, and chemotherapy involving mitomycin C (MMC) and/or cisplatin. For more advanced NMIBC, the second-line chemotherapeutic gemcitabine can also be used in combination with cisplatin or another platin drug such as paclitaxel (9). The chance of recurrence of high-risk NMIBC five years after TURBT is approximately 69.5%, with up to 22.8% progressing to MIBC (10).

First-line treatment for MIBC includes neoadjuvant chemotherapy followed by radical cystectomy, i.e. complete bladder removal and the implementation of a stoma. The impact on such patients can be significant, with overall survival post-treatment at 49% after 5 years and 36% at 10 years (11). In addition, those who survive nevertheless suffer from major quality-of-life issues involving social and psychological impacts (12, 13). This highlights the importance of developing new therapies and models to test them, especially at the NMIBC stage before progression has occurred (14).

The most widely used preclinical models for bladder cancer are orthotopic and xenograft murine models. However, working with mice is labour-intensive, ethically challenging, expensive and subject to time-consuming regulation. Other disadvantages include the length of time needed to establish disease in mice, which can take 4-6 months, and in the case of the PDX model, poor implantation rates (15). More profoundly, there are concerns about physiological inaccuracies caused by the species difference of the surrounding urothelial environment (16). It has been shown that the method of tumorigenesis and the tumour microenvironment greatly influence the molecular subtype which consequently influences cancer development and response to treatment (17). Therefore, in the mouse model context, even implanted human bladder cancer cells may not respond as they would in a human context. This highlights the need for developing complementary approaches to model bladder cancer.

A more representative model would involve combining human tumour cells with a human microenvironment. On the cancer side, 3D spheroid models provide more complexity than simple tumour cell lines, allowing cells to communicate with one another and express chemical gradients shown in normal tumour progression such as nutrients, oxygen and catabolites (18). Tumour spheroids can also be derived from patients, which may usher in an era of personalized medicine for bladder cancer (19).

On the microenvironment side, recent years have seen exciting advances in healthy human urothelial organoid and microtissue modelling (reviewed in (16, 20)). Briefly, human cell lines and patient-derived cells have been used including UROtsa (21) human embryonic stem cells (hESC) (22) and normal human urothelial cells (NHU) (23, 24). Cross *et al*. developed a protocol allowing primary NHU cells to stratify to 3-7 layers with expression of a number of key markers (25). More recently, Sharma et al. developed a human bladder-on-chip model for studying UTI with UPEC infection. Human bladder endothelial and epithelial cells were co- cultured under flow conditions using highly diluted urine. Significantly, the model also incorporated neutrophils as an immune component (26).

Recently, our laboratory has developed a <u>3D</u> <u>u</u>rine-tolerant <u>h</u>uman <u>u</u>rothelium model (3D-UHU) which has previously been characterised and used for research into urinary tract infections (27, 28). This microtissue model, derived from commercially available HBLAK cells, is novel because it can be grown and maintained in a 100% urine environment. Cells stratify to 5-7 cell layers (as opposed to the mouse urothelium, which only expresses three) with terminally differentiated umbrella cells. Urothelial biomarkers are also correctly expressed (Claudins, Uroplakins, Zonula Occludens-1 (ZO-1), Cytokeratins, Toll-like receptors (TLRs), E-Cadherin) with the formation of a luminal glycosaminoglycan layer and strong barrier function. Importantly, it is fully urine-tolerant for weeks, allowing for experimentation in the native environment, which may affect both tumour and normal urothelial gene expression as well as response to chemotherapy, providing a more physiologically relevant tumour microenvironment for understanding both basic biology and treatment response.

Here, we created and characterised hybrid tumour/healthy cell models using the 3D-UHU model co-cultured with bladder cancer spheroids derived from bladder carcinoma cell lines frequently used in mouse models, RT112 (low grade) and T24 (high grade). Next, we characterised the models with various markers involved in diagnosis (CK7, CK20, GATA3), adhesion (cadherins), invasion/migration (MMPs), and proliferation (Ki-67). Finally, we assessed the usefulness of 3D-UHU-TU as a testbed for treatment using the common chemotherapeutic Mitomycin C. Our results suggest that the models hold promise for trialling potential therapies in a manner far simpler, quicker and cheaper than animal experimentation. It also allows for testing in a human urinary microenvironment, where urothelial side effects can be assessed alongside tumour cell killing.

## 2. Methods and Materials

### 2.1 Cell Lines and Culture Conditions

Human HBLAK cells (CELLnTEC, Switzerland) were cultured as described previously (27). Briefly, HBLAK cells were cultured in T150 flasks in 2D Prime media (CnT-PR; CELLnTEC, Switzerland) at a seeding density of 3x10^5^. Cells were incubated at 37°C at 5% CO2 for 5-6 days with media changed every 2-3 days until 80-90% confluent. It was important that the cells did not exceed 90% confluency to avoid early cell differentiation. Cells were detached using Accutase solution (CELLnTEC, Switzerland) and collected by centrifugation at 300 x g for 5 mins. The cells were either passaged up to P13 or used for 3D cell culture.

The RT112 (low-grade) human bladder cancer cell line was kindly donated by Nenna Kanu and Dr Mark Linch (University College London, UK). The T24 (high-grade) bladder cancer cell line was purchased from ATCC. RT112 and T24 were cultured in complete DMEM (Thermo Fisher Scientific, UK) with 10% foetal bovine serum (FBS; Thermo Fisher Scientific, UK) supplemented with Pen/Strep solution (100U/ml; Merck, UK; A5955). All cells were incubated at 37°C with 5% CO2. Cells were harvested at ∼90% confluency and detached with 0.25% Trypsin-EDTA (Thermo Fisher Scientific, UK) and collected by centrifugation at 300 x g for 5 mins. Cells were passaged or used for spheroid generation.

### 2.2 3D-UHU Cell Culture

3D-UHU microtissue models were created as described previously (27, 28). Briefly, at ∼90% confluency, HBLAK cells were harvested between passages 8-13 with Accutase solution and added to 0.4µm polycarbonate Transwell membrane inserts (Corning, USA) at a seeding density of 4x10^5^ in a total of 400 µl 2D Prime media in the apical chambers. 1.5ml 2D prime media was added to the basal chambers. Transwell membranes were 100% confluent after 48 hours, and pre-warmed 3D Prime media (CnT-PR-3D; CELLnTEC, Switzerland) was added to apical and basal chambers to initiate stratification and differentiation. After at least 72 hours’ incubation, Transwells were transferred to 12-well Thincert® plates (Greiner Bio-One, UK), and 300 µl of filtered pooled-gender human urine (BioIVT, UK) from healthy donors were added to the apical chambers and 4.3 ml 3D Prime media was added to the basal chamber. Media and urine changes were performed every 3-4 days for 14-18 days. 3D-UHU organoids were either used for further experiments or processed for immunofluorescence imaging (see below).

### 2.3 RT112/T24 Spheroid Generation and Growth Curves

Trypsinised RT112 and T24 cells were collected (see above) and resuspended in complete culture media separately. CellTracker Green CMFDA dye (Invitrogen, UK) was then added to an aliquot of centrifuged cells according to the manufacturer’s instructions. Dyed cells were seeded into Elplasia microspheroid plates (Corning, UK) at a density of 1,000 cells per microcavity (79,000 cells /well) in 200µl volume of media and incubated at 37°C with 5% CO2 for 2-4 days. For growth curves, diameters of spheroids were measured each day for 6 days (n=12). Spheroids were imaged using a Leica DMi1 inverted microscope and captured using LAS X software V5.0.2.24429. Images were analysed using Image J V1.54f (29).

### 2.4 3D-UHU-TU Establishment: Incorporation of Cancer Spheroid onto 3D-UHU model

To establish 3D-UHU-TU models (Figure 1), HBLAK cells were grown in 3D conditions for at least 14-18 days as described above. On the day of spheroid implantation, a full media/urine change was performed on the 3D-UHU models. Spheroids were then harvested using a wide-bore 200µl tip. 150 µl were aspirated up and down gently to dislodge spheroids from the microcavities and then collected. The spheroid suspension was next deposited into the urine slowly and at different positions on the Transwell to disperse the spheroids evenly. The 3D- UHU-TU models were then incubated overnight at 37°C. The next day, the urine/DMEM solution was changed to fresh urine to ensure a 100% apical urine environment. 3D-UHU-TU models were then used for MMC treatment experiments or analysed with immunofluorescence staining (see below).

**Figure 1:**
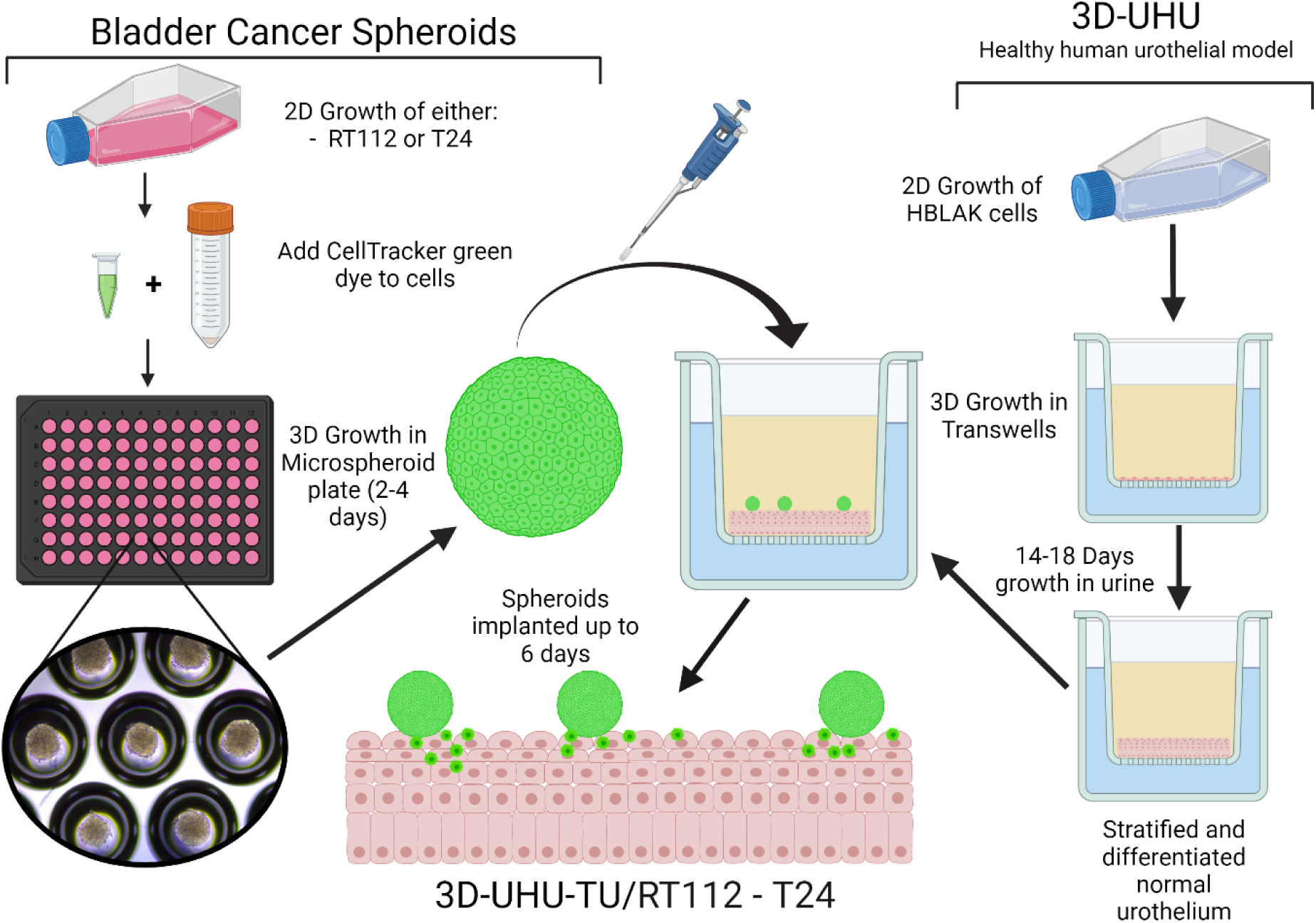
Schematic workflow for generating the 3D-UHU-TU model. Images not to scale. Created with BioRender.com.

### 2.5 Immunofluorescence Staining and Antibodies

All samples in the Transwells were fixed in 4% methanol-free formaldehyde (PFA; Fisher Scientific, UK) in phosphate-buffered solution (PBS; Thermo Fisher Scientific, UK) for 4°C overnight. Transwell membranes were cut out using a scalpel and placed in a well of a 24-well plate. 0.2% Triton-X100 (Sigma Aldrich, UK) was added to each sample to permeabilise cells for 35 mins under cover at room temperature with gentle rocking at 15 rpm. Triton-X100 was aspirated out and samples were washed once with PBS.

For antibody staining, blocking solution comprised of 5% normal goat serum (NGS; Thermo Fisher Scientific, UK) with 1% bovine serum albumin (BSA; Merck, UK) in PBS was added to the samples. These were covered for 1 hour at room temperature with gentle rocking at 15 rpm. Blocking solution was aspirated and samples were washed once with PBS with 1% BSA. Primary antibodies (Table S1) were suspended in 1% NGS and 1% BSA in PBS, added to samples, and incubated at 4°C overnight. Next, samples were washed three times with 1% BSA in PBS. Secondary antibodies (Table S1) were suspended in 1% NGS and 1% BSA in PBS and added to wells under cover for 1 hour at room temperature with gentle rocking at 15 rpm. Secondary antibodies were aspirated and samples were washed with PBS three times.

4′,6-diamidino-2-phenylindole (DAPI; Thermo Fisher Scientific, UK) and Phalloidin-555 or 647 (Thermo Fisher Scientific, UK) (Table S1) were diluted together in PBS and added to samples and incubated for 1 hour under cover at room temperature with gentle rocking at 15 rpm. The DAPI and Phalloidin solution was aspirated and samples were washed 4 times with PBS. All samples were transferred onto glass slides and mounted using Prolong Diamond antifade mounting solution (Invitrogen, UK). Coverslips were placed on the samples with the edges sealed with nail varnish and allowed to dry overnight at room temperature before imaging.

Samples were then imaged using Leica SP8 confocal microscope and analysed in Image J V1.53t (29).

### 2.6 Immunohistochemistry

3D-UHU-TU models were fixed in 10% neutral buffered formalin for no longer than 24 hours and then inked with 0.1% safranin O and bisected. These were processed using the Leica Peloris II Tissue Processor (Leica Biosystems) and embedded on edge into paraffin blocks using the Tissue-Tek TEC 6 embedding centre (Sakura Finetek USA) with the cut surface of the bisected wells facing down. 3 µm sections were obtained through Leica Histocore Autocut microtome (RM225, Leica Biosystems), mounted on TOMO adhesive slides (SolMedia) and baked in a 60°C oven before staining with haematoxylin and eosin (H&E) on the Leica Autostainer X (Leica Biosystems). Immunohistochemistry (IHC) staining with GATA3 antibody (L50-823, Cell Marque, Sigma Aldrich), CK7 (RN7, Leica), and CK20 (SP33, Roche) was performed on the Leica Bond Max (Leica Biosystems) and Ventana Benchmark Ultra (Roche) automated immunostaining platforms. GATA3 antibody was diluted with a primary antibody diluent (AR9352, Leica Bond, Leica Biosystems) 1:300 prior to use; the other antibodies mentioned were used undiluted.

### 2.7 Mitomycin C Treatment of 3D-UHU-TU Models and Components

3D-UHU-TU models were established as described above. Mitomycin C (MMC) (11435-25mg- CAY; Cambridge bioscience, UK) stocks were diluted in pre-warmed 0.85% saline (Fisher Scientific, UK) to 10 and 50 µg/ml working solutions. Working MMC solutions were added to the 3D-UHU-TU models in duplicate and incubated for 2 hours at 37°C. After treatment, the MMC solution was aspirated out and fresh urine was added. Treated 3D-UHU-TU models were incubated for 24 hours before being fixed, stained and imaged with the SP8 confocal microscope as described above.

### 2.8 Image Analysis and Graph Generation

3D-UHU-TU models were analysed using ImageJ V1.53t-1.54f and spheroid growth curves were generated using GraphPad Prism V9.1.2.

## 3. Results

### 3.1 Establishment of Cancer Spheroids and 3D-UHU-TU Model

To create a model combining the human tumour cells into the healthy 3D-UHU microenvironment (Figure 1), we chose human bladder cancer cells lines RT112 (low grade), and T24 (high grade) for tumour spheroid generation. These cell lines have been frequently used in orthotopic mouse models (30), making our data comparable with the current literature. RT112 and T24 cells were seeded into ultra-low attachment microwell plates and grown in complete DMEM for 2-4 days; the resulting spheroids reached an average diameter of 200µm and 140µm respectively (Figure 2).

**Figure 2:**
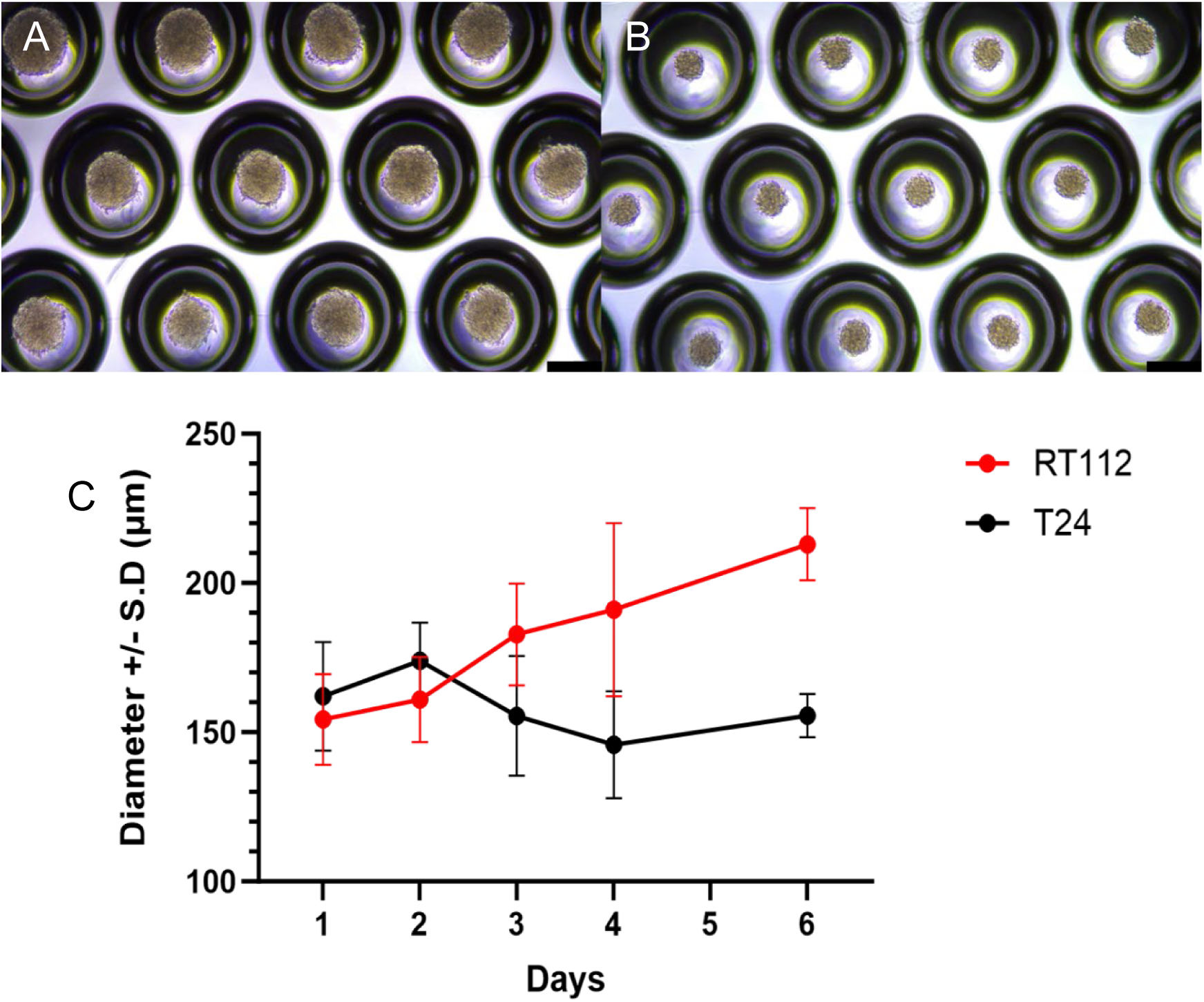
RT112 and T24 spheroids in Elplasia microcavity ULA plates, grown in standard DMEM. (A) Day 4 RT112 spheroids ; (B) Day 4 T24 spheroids ;(C) Growth curves of RT112 and T24 spheroids over 6 days. Growth curves generated from 12 technical replicates from 1 biological experiment. Scale bar = 200µm.

3D-UHU models contained 5-7 layers of urothelial cells, and all had a differentiated umbrella cell layer (Data not shown). When incorporated onto the 3D-UHU surface and maintained in 100% urine for at least 24 hours, RT112 and T24 spheroids exhibited different phenotypes. RT112 spheroids were larger and maintained their spherical shape when integrated onto a stratified 3D-UHU model (Figure 3A; note that in the confocal image in 3A, the coverslip has flattened the spheroid somewhat. In Figure S1, RT112 models captured using Light Sheet microscopy show the full spherical shape). In contrast, T24 spheroids readily spread out over the surface of the urothelium with the spheroid edges migrating away from the main spheroid mass and integrating extensively into the top layer of the urothelium (Figure 3B). 3D-UHU-TU created with both high- and low-grade cell lines were morphologically distinct.

**Figure 3:**
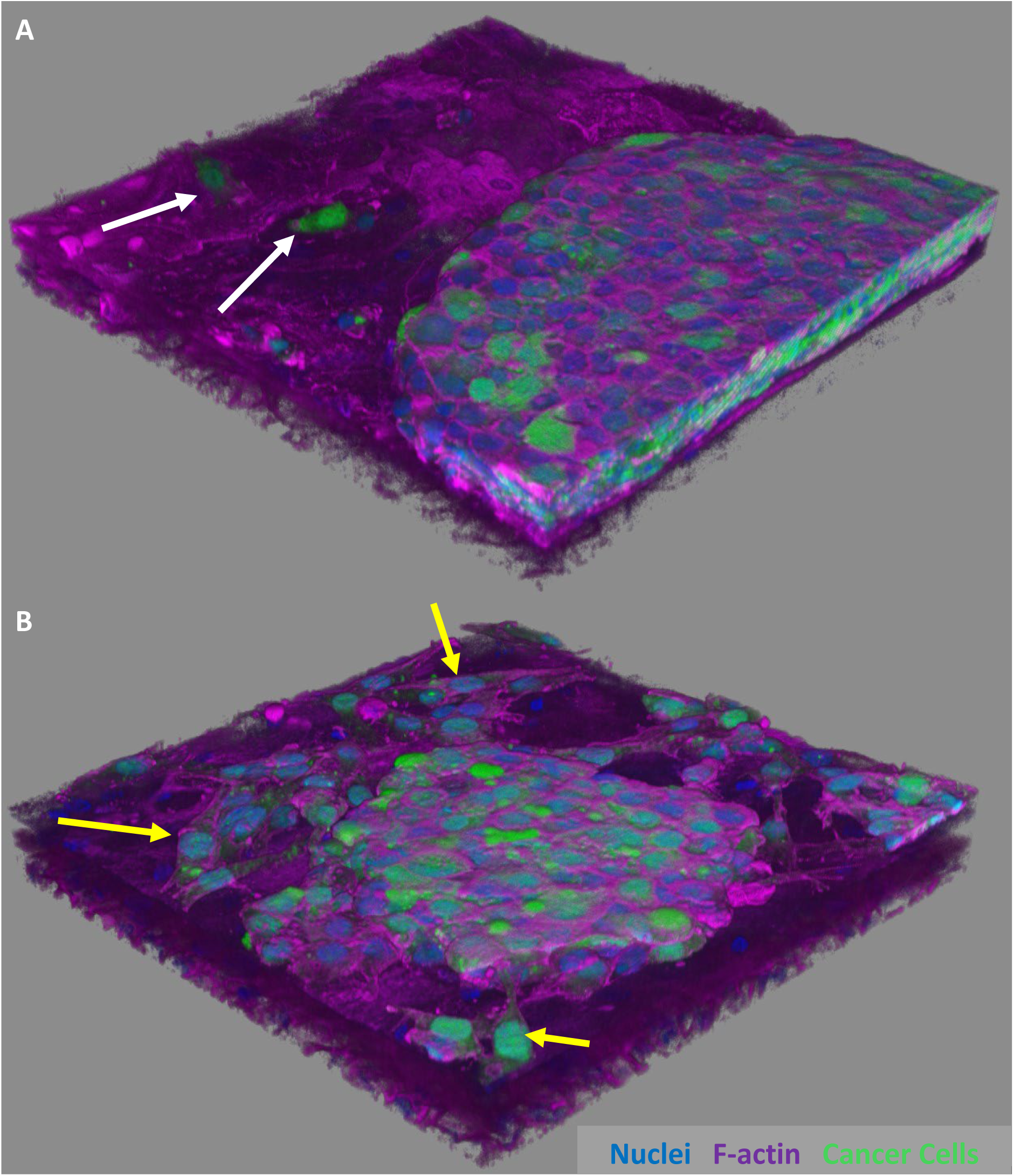
Optimised 3D-UHU-TU with RT112 spheroids and T24 spheroids tagged with CellTracker CMFDA probes (green). (A) RT112 spheroids flattened and maintained their spherical shape with some migration on top of the urothelial surface. Migrating cells appeared to integrate into the urothelium (white arrows) (B) T24 spheroids were smaller and exhibited more migration of cells at the spheroid edge (yellow arrows). Representative images from 3 independent experiments.

### 3.2 Characteristics of 3D-UHU-TU Models via Immunohistochemistry and Immunofluorescence

#### 3.2.1 Diagnostic Biomarkers CK7, CK20, GATA3

To determine whether the 3D-UHU-TU models retained spatial markers reminiscent of patient-derived bladder cancer tissue, we first used immunohistochemistry to assess expression of several key proteins. In addition, to inspect general morphology of the 3D-UHU- TU models maintained in 100% urine environment for 6 days, we performed haematoxylin and eosin (H&E) staining on fixed, paraffin-embedded sections, orthogonally bisecting the area of the microtissue containing a spheroid.

The H&E stain shows the compact structure of RT112 spheroid (Figure 4A) which matches the previous images (Figure 3A, Figure S1). Similarly, the cell migration from the T24 spheroid was consistent with the metastatic features of this high-grade cell line in both IF and H&E (Figure 3B and 4A respectively).

**Figure 4:**
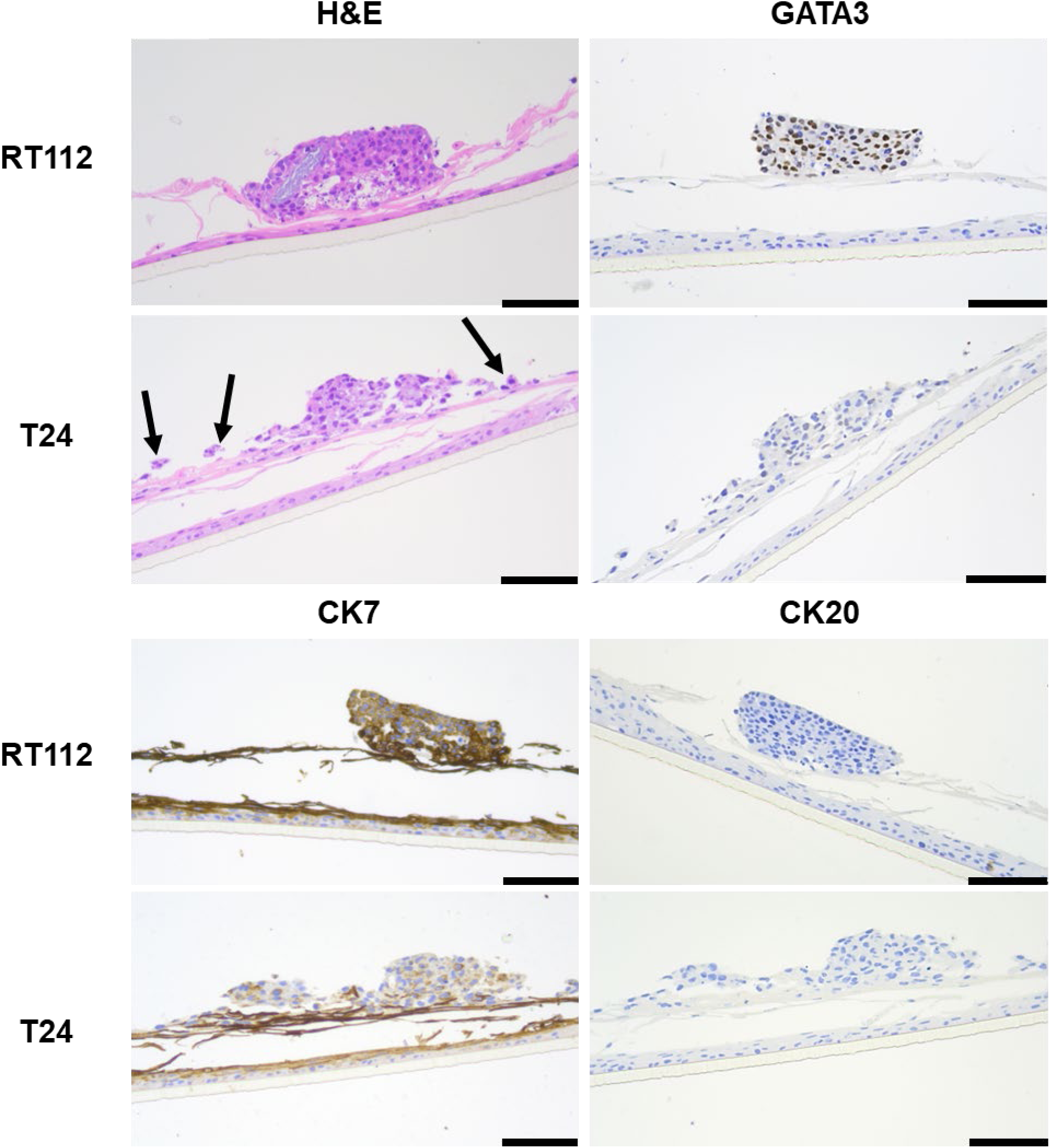
Immunohistochemistry and H&E stain of 3D-UHU-TU models. (A) H&E stain of both 3D-UHU-TU models. Expression of diagnostic markers (B) GATA3; (C) CK7 and (D) CK20. 3D-UHU-TU models maintained up to 6 days in 100% urine environment. Images are representative of 3 separate biological replicates. Scale bar = 50µm.

Next, we inspected markers commonly used in bladder cancer diagnosis. GATA3 is a transcription factor typically found in the nuclei of lower-grade bladder cancer but is down- regulated in higher-grade cancers, making it an important diagnostic marker for tumour grading (31). In the 3D-UHU-TU models, GATA3 was expressed in the low-grade RT112 cancer spheroid but not in the high-grade T24 cancer spheroid as expected (Figure 4B).

Expression of CK7 and CK20 is commonly used in the diagnostic laboratory. CK7 is expected to be expressed in all layers of the urothelium and in cancer. Indeed, CK7 was strongly expressed in both the cancer spheroid and healthy urothelial components in the model (Figure 4C). CK20 expression is known to be decreased in bladder cancer tissue (32), and is known to be expressed on the apical surface of the healthy urothelium (33) and of the 3D- UHU model (21). In both RT112 and T24 models, CK20 was not detected in either the tumour or the surrounding urothelium (Figure 4D).

#### 3.2.2 E- and N-Cadherin Expression

Cadherins are cell-to-cell adhesion mediators that are expressed in bladder cancer and are a known indicator of cancer progression (34). E-Cadherin is expressed in healthy epithelium and Ta tumours, but switch to N-Cadherin in more invasive, high-grade tumours (35). E- and N-Cadherin were examined for correct expression in both high -and low-grade models (Figure 5; 3D-UHU not shown). As expected, E-Cadherin was expressed in the low-grade RT112 spheroids (Figure 5A), localised correctly at cell-cell junctions. It was also expressed in the healthy urothelial component, particularly in the superficial umbrella cell layer (Figure 5C) and similarly enriched at junctions. In the T24 spheroids, as expected for a high-grade cancer, E-Cadherin was scarcely apparent, with possible faint expression on the edges of the spheroid. Reciprocally, N-Cadherin was not expressed in the RT112 spheroids, but was in T24 spheroids (Figure 5B), both also as expected.

**Figure 5:**
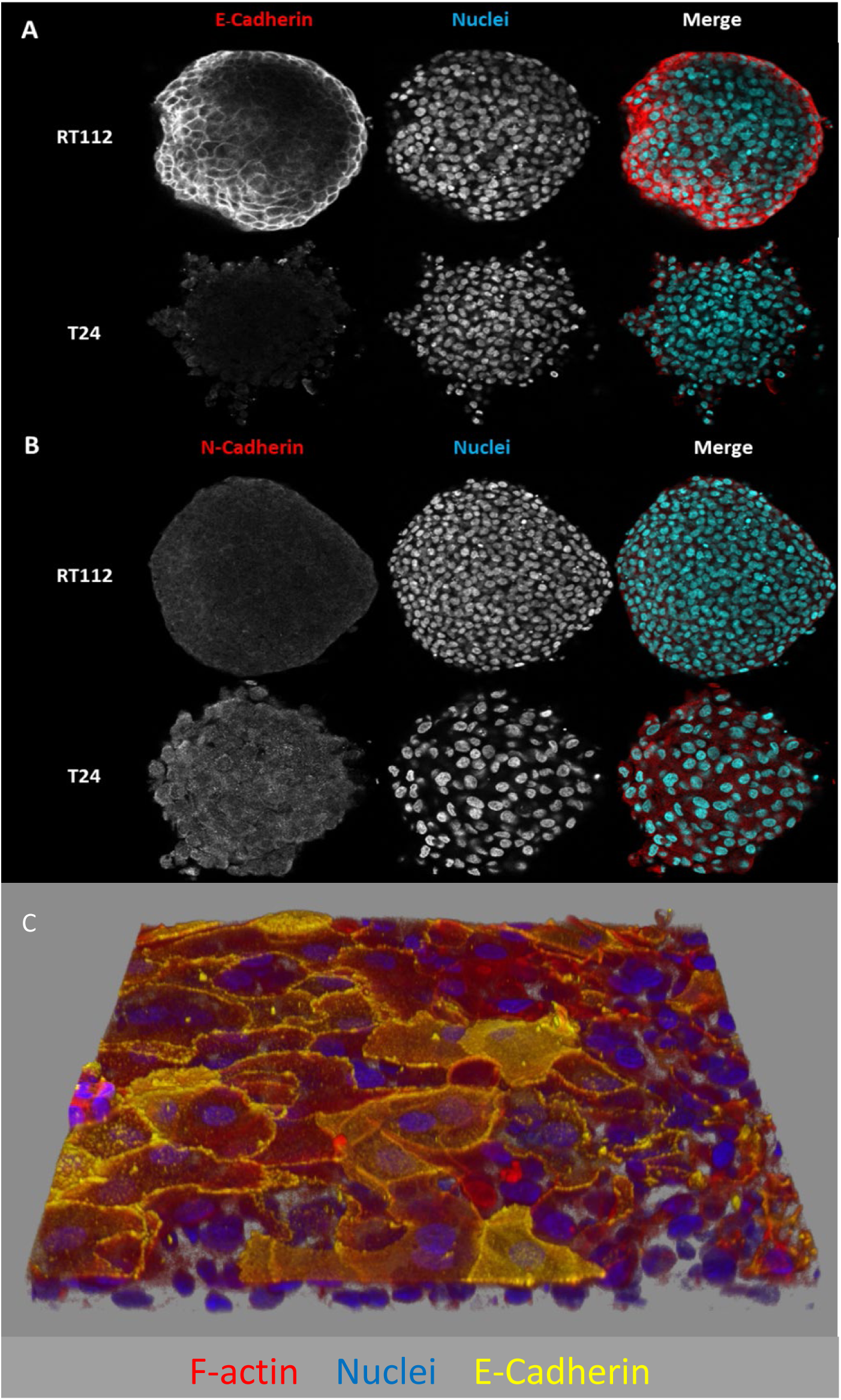
Expression of E- and N-Cadherin in RT112 (low grade) and T24 (high grade) cancer spheroids and 3D- UHU in 100% urine environment. (A) cancer spheroids were co-cultured with optimised 3D-UHU model (3D-UHU layer is below the plane of focus and not apparent here) with expected expression of E-Cadherin in RT112 spheroids and expression of N-Cadherin in T24 spheroids. (B) 3D-UHU model expressed E-Cadherin in differentiated umbrella cells as expected. Images are representative of 3 independent experiments.

#### 3.2.3 Invasion and Migration Marker Matrix Metalloproteinases -2 and -9

Matrix Metalloproteinase (MMP) -2 and -9 (also known as gelatinase A and B respectively) are important for cancer progression particularly for immunomodulation, cell migration, invasion and morphogenesis of different cancers (36). We wanted to examine expression of these metalloproteinases on the cancer spheroids integrated onto 3D-UHU to determine whether these hallmarks of cancer progression could be obtained in an *in vitro* 100% urine environment. In particular, these markers help assess cell migration and potentially invasion.

MMP-2 and MMP-9 were expressed on both RT112 and T24 cancer spheroids (Figure 6A&B; 3D-UHU component present but not shown). In RT112 cancer spheroids, MMP-2 and -9 are expressed in the main spheroid but also in cells that migrate out of the main spheroid (white and yellow arrows; Figure 6A and B). These cells also seem to form a ‘chain-like’ structure (blue arrow; Figure 6B) and have a mix of amoeboid (white arrows; Figure 6A) and mesenchymal (yellow arrow; Figure 6B) migration phenotypes.

**Figure 6:**
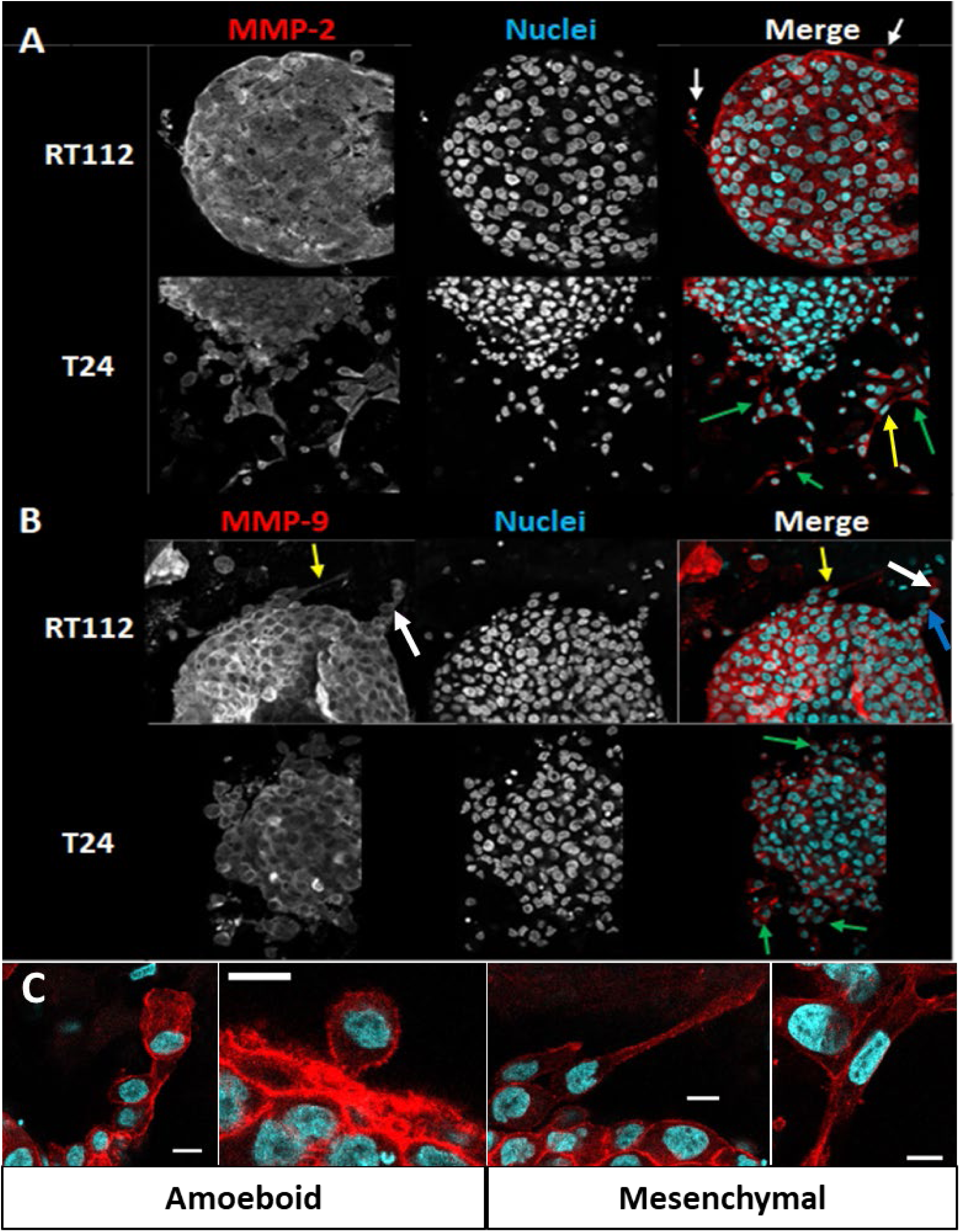
MMP-2 (A) and MMP-9 (B) expression of RT112 and T24 spheroids integrated onto the 3D-UHU model (3D-UHU layer is below the plane of focus). MMP-2 and -9 (Red, Panels A and B) are expressed in the correct areas with different migration phenotypes of cells migrating from the main spheroid. In RT112 spheroids, ‘chain-like’ structures are present (blue arrow) with migration phenotypes include amoeboid (white arrows) and mesenchymal (yellow arrows). In T24 spheroids, there were more migrating cells (green arrows) as expected of a higher-grade cancer. There was also a mesenchymal migration phenotype present (yellow arrow). (C) Migration phenotypes observed in 3D-UHU-TU models. Amoeboid phenotype found in RT112 spheroids (both panels) and mesenchymal found in both RT112 and T24 spheroids (Left panel = RT112; Right panel = T24). Red = F-actin (Panel C only). Scale bars = 10µm. Representative images from N=3 independent experiments.

Similarly, in the case of T24 spheroid and migrating cells, both expressed MMP-2 and -9. There were a lot more cells that had migrated out of the main spheroid (green arrows; Figure 6A&B compared with RT112 spheroids, which is expected of a higher-grade cancer. These cells also only expressed mesenchymal migration phenotype (yellow arrow; Figure 6A).

#### 3.2.4 Proliferation – Ki-67

Ki-67 is a widely used proliferation marker in cancer cell biology. It is a vital protein of the cell cycle which is upregulated in proliferating cells and downregulated in resting cells (37). It is also a promising prognostic marker particularly for breast cancer, indicating its clinical relevance (38).

In the low-grade RT112 model, the initial experiment of co-culture for 2 days in a urine environment showed expression of Ki-67 expression in the RT112 spheroid, and also in migrating cells (Figure 7A). This was also confirmed with histological analysis after 6 days of co-culture in urine (Figure 7B). These results indicate that these cancer spheroids and migrating cells could survive and proliferate in a urine environment.

**Figure 7:**
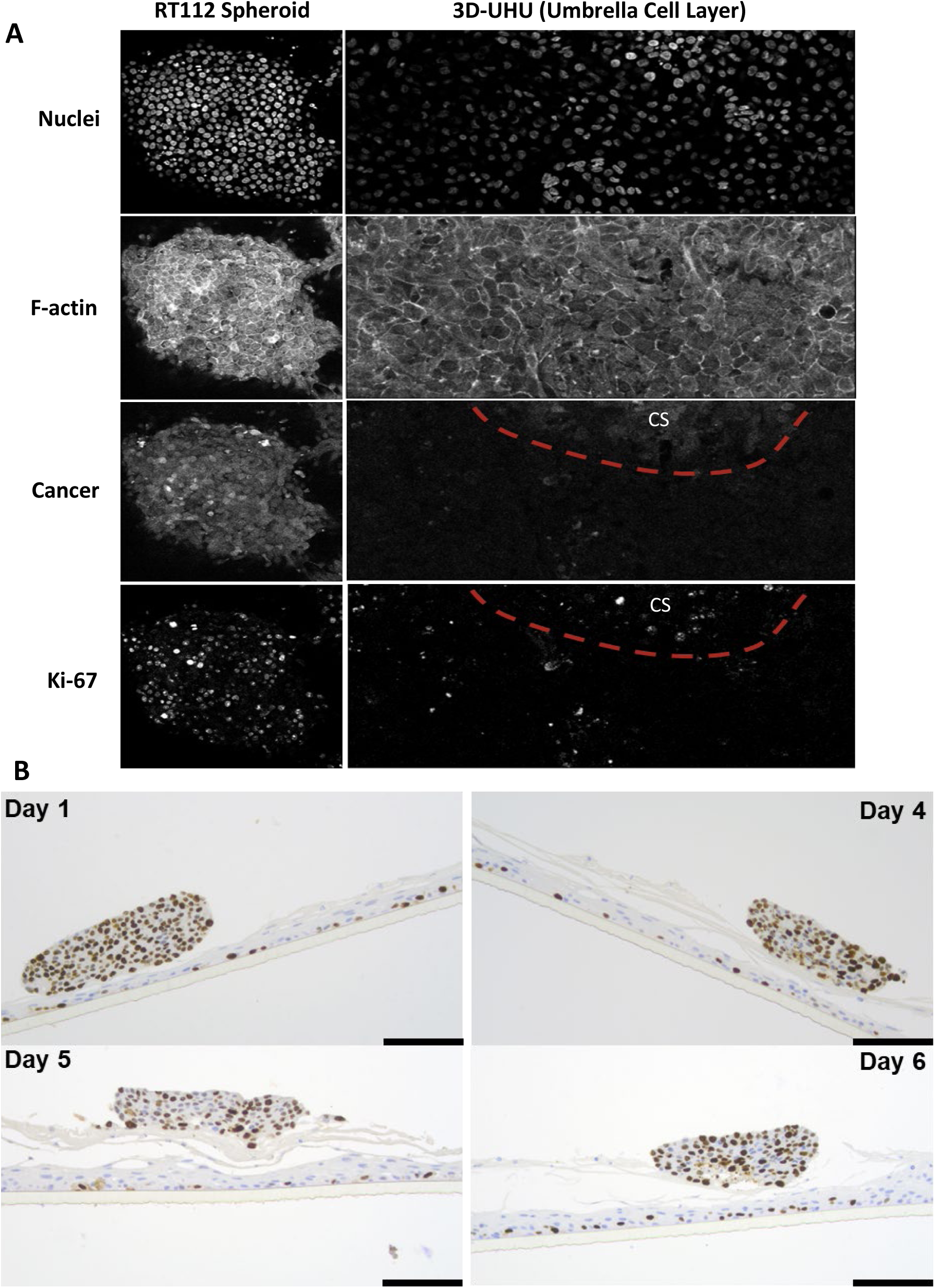
Expression of Ki-67 proliferation marker in 3D-UHU-TU/RT112. (A) Split channels of immunofluorescence with separate RT112 spheroid and 3D-UHU after 4 days in co-culture. ‘CS’ indicates cancer spheroid. (B) Immunohistochemistry staining of Ki-67 on 3D-UHU-TU/RT112 maintained up to 6 days in 100% urine environment. Positive expression is seen in the main cancer spheroid and basal layer of the urothelium. Scale bar = 50µm. Images are representative of 3 independent experiments.

In contrast, the high-grade T24 model exhibited limited Ki-67 expression, particularly within the T24 cancer spheroids (Figure 8A and B; right of the orange dotted line). However, while T24 cells proliferated normally in 2D DMEM cultures, they appeared to struggle in urine conditions, particularly within the 3D spheroid structure. In contrast, T24 cells migrating away from the spheroid mass expressed Ki-67 (Figure 8A).

**Figure 8:**
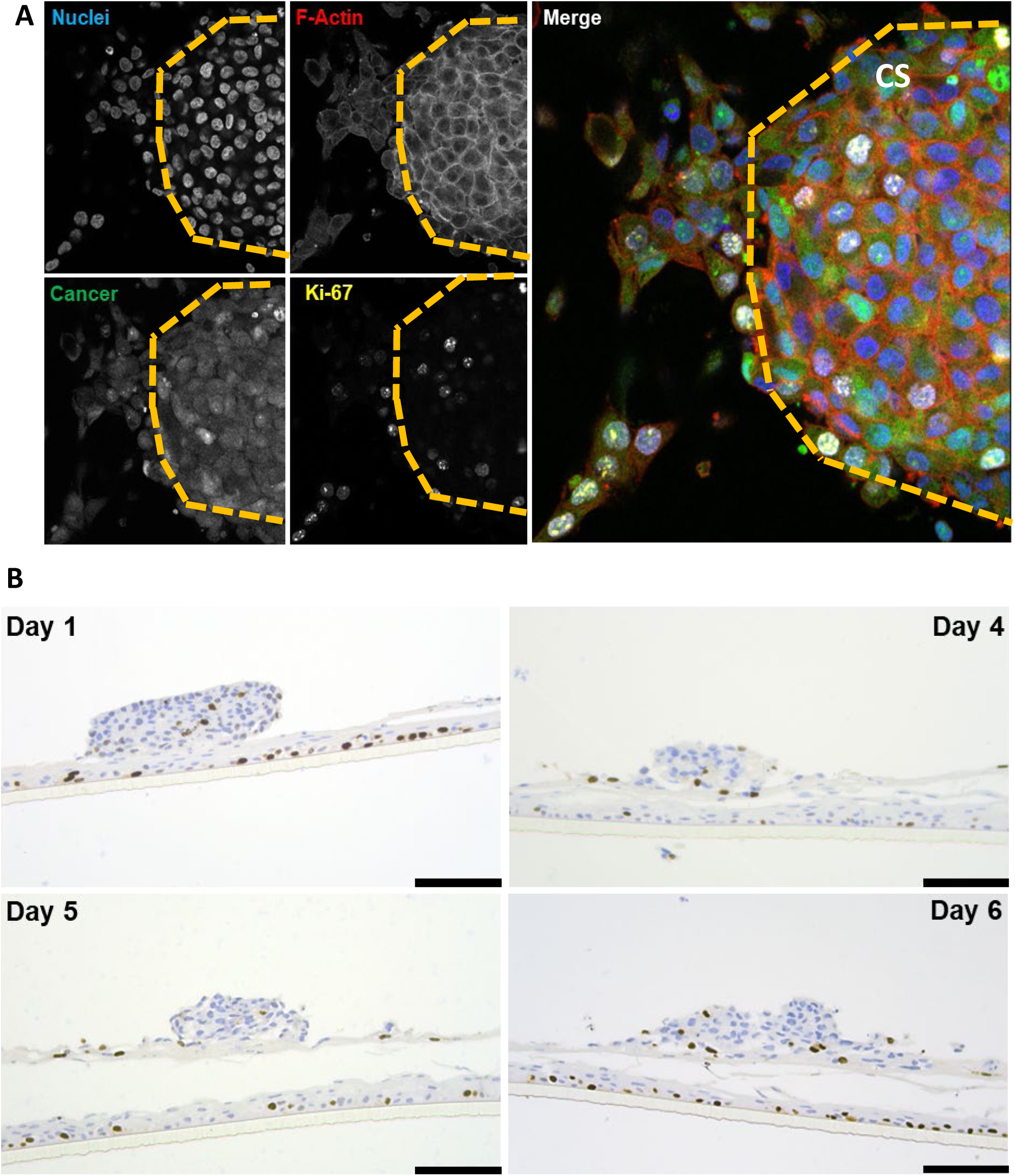
Expression of Ki-67 proliferation marker in 3D-UHU-TU/T24. (A) Split channels of immunofluorescence with separate T24 spheroid and 3D-UHU after 2 days in co-culture. Cancer spheroid (CS) is located to the right of the orange dotted line. Some expression in migrating cells but not in the main spheroid. (B) Immunohistochemistry staining of Ki-67 on 3D-UHU-TU/T24 maintained up to 6 days in 100% urine environment. Minimal expression is seen in the main cancer spheroid but is positive in the basal layer of the urothelium. Scale bar = 50µm. Images are representative of 3 independent experiments.

Histological analysis of the models further revealed Ki-67 expression in the basal cells of the surrounding normal urothelium, as well as in some intermediate cells. This is expected in the transitional urothelium, and the presence of Ki-67 in these layers supports the validity of the healthy component of the 3D-UHU-TU model.

### 3.3 Mitomycin C Treatment has Tumour-Specific Efficacy on 3D-UHU-TU Models

Mitomycin C (MMC) is one of the most commonly used chemotherapeutic agent for treatment of NMIBC (39). It works by alkylating cellular DNA, causing DNA double-strand breaks that initiate apoptosis (40). We trialled this gold-standard MMC treatment in the 3D- UHU-TU models to validate the platform as a potential testbed for new treatments. MMC treatment was administered, and specific cytotoxicity assessed via immunofluorescence using confocal microscopy. Two treatment were used, a low-dose (10μg/ml) and a high dose (50µg/ml), for two hours; afterwards, the treatment was replaced with urine and models were allowed to recover for 24 hours before assessment.

For the RT112 models (Figure 9; left panels), MMC caused damage to the RT112 spheroids in a dose-dependent manner, as expected. While the saline control did not induce morphological damage or affect the spheroid structure, as expected, (Figure 9; top-left panel), at 10µg/ml, the structure of the spheroid was still intact, but extensive cellular lysis and nuclear damage was evident (Figure 9; middle-left panel, white arrows). At 50µg/ml, the RT112 spheroid appeared to be completely disaggregated with the healthy urothelium still intact (Figure 9; bottom left panel).

**Figure 9:**
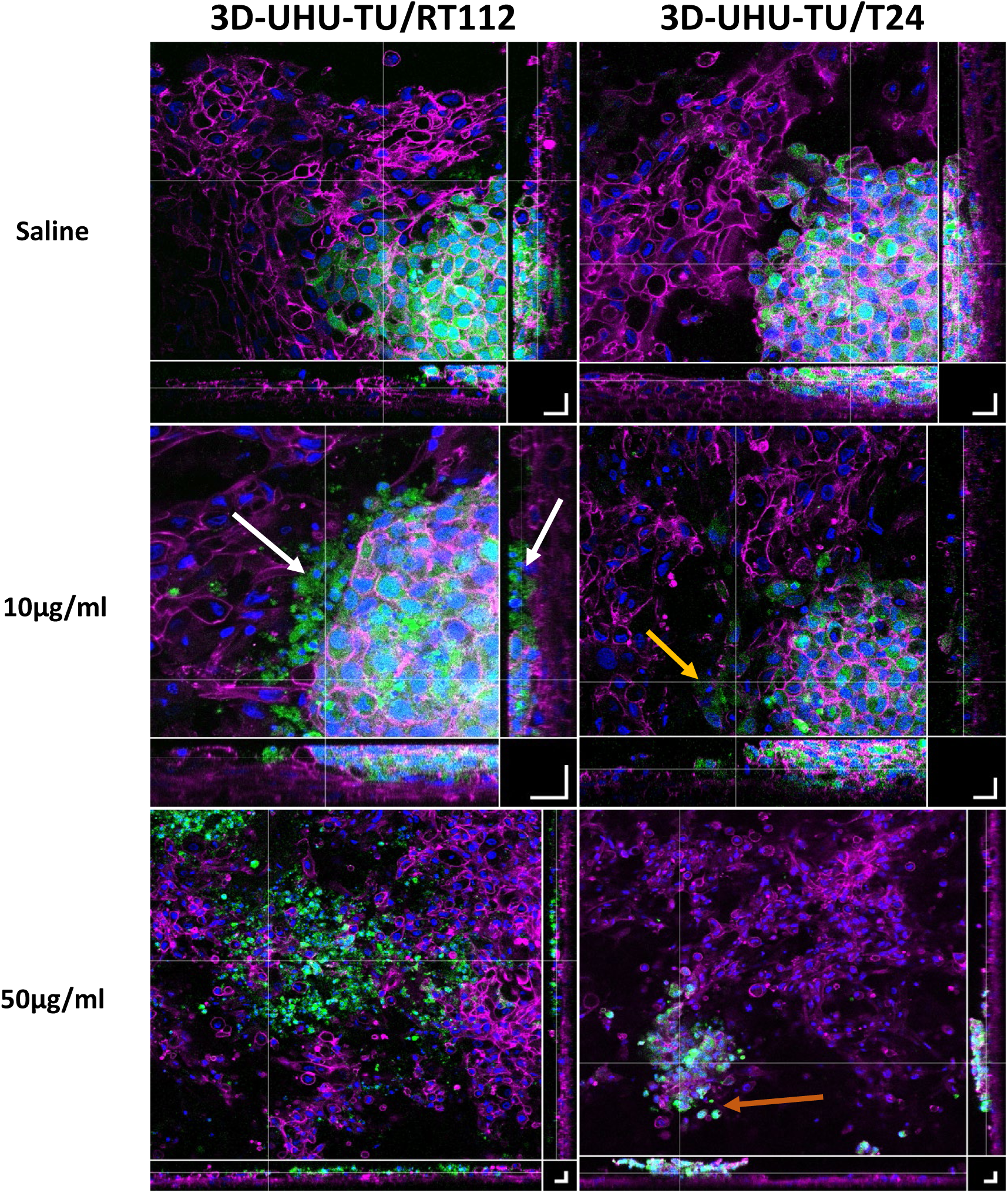
MMC treatment of 3D-UHU-TU models. The spheroid component of the low-grade RT112 models (left panels) lose spheroid structure and exhibit cellular lysis and damage (white arrows) when treated with a lower dose of 10µg/ml (middle left). Treatment with 50µg/ml (bottom left) showed complete disaggregation of the cancer spheroid with urothelial layer still intact. High-grade T24 models (right panels) also showed cellular damage of migrating cells (orange arrow) when treated with 10µg/ml MMC (middle right). At 50µg/ml (bottom right), the cancer spheroid was much smaller and more cellular damage is observed (brown arrow). Scale bars = 20µm. Images are representative of 3 independent experiments.

In the T24 models (Figure 9; right panels), saline treatment again induced no morphological damage to the model (Figure 9; top-right panel). At 10µg/ml MMC, the cancer spheroid appeared to maintain structure with some cellular damage of migrating cells, comparable to what was observed with RT112 (Figure 9; middle-right panel, orange arrow). At 50µg/ml, the cancer spheroids were much smaller and some cellular damage was observed (Figure 9; bottom-right panel, brown arrow).

## 4. Discussion

The 3D-UHU model has previously been utilised for research into urinary tract infection (27, 28). 3D-UHU-TU represents the first significant advancement in adapting this model to simulate another urological disease: bladder cancer. Not only is the 3D-UHU-TU model established in a healthy urothelial and urine-dependent environment, allowing for a more physiologically accurate tumour microenvironment, but it can also incorporate tumour spheroids from different origins – in this case, both low- and high-grade 3D bladder cancer spheroids were represented as a proof-of-principle. Characterisation of the 3D-UHU-TU models revealed them to largely express the expected markers in the correct spatial environment. It is worth mentioning that growth and implantation of larger spheroids (≥400µm) is also feasible (Figure S1), so there is also scope to assess larger spheroids that incorporate a hypoxic and/or necrotic core (41). However, due to imaging scale constraints, to obtain more accurate spatial information, we focused on the implantation of multiple RT112 and T24 micro-spheroids in this paper.

Clinical diagnoses of bladder cancer using immunohistochemistry staining usually involve GATA3 (42), CK7 and CK20 (43) and these three biomarkers were chosen to assess whether the 3D-UHU-TU models expressed similar diagnostic markers to patient material. Firstly, the loss of GATA3 expression has been linked to higher-grade disease (44). A tissue microarray study of 2,710 bladder tumours found GATA3 to be significantly more prevalent in non-invasive Ta stage tumours (98% of 790 tumours), with a marked reduction in T2 to T4 stage tumours (59.8% of 1,223 tumours) (31). Since RT112 is a low-grade T2-stage cancer cell line, we anticipated GATA3 expression, and indeed, the 3D-UHU-TU/RT112 model demonstrated GATA3 expression in the spheroid component. In contrast, the 3D-UHU-TU/T24 model, consistent with the characteristics of a high-grade cancer cell line, did not exhibit GATA3 expression in the spheroid component.

CK7 and CK20 expression were analysed in another microtissue array study in various grades and stages of urothelial carcinoma across 1,755 samples (45). In Ta stage and low-grade urothelial carcinoma, CK7 and CK20 were expressed in 98.7% and 69.7% of 177 tumours, respectively. In the 3D-UHU-TU/RT112 model, which mirrors these types of tumours, CK7 was consistently expressed throughout the model. However, there was little expression of CK20 in the spheroid mass. Like the RT112 model, the 3D-UHU-TU/T24 model also expressed CK7, though at a weaker level in the cancer spheroid compared with the RT112 cancer spheroid. CK20 was also not expressed in the T24 spheroids.

Although CK20 has previously been shown to be expressed in the superficial umbrella cells of 3D-UHU by immunofluorescence analysis (27), we did not observe CK20 expression via ICH in the normal margins of any of our models. Positive staining controls were employed CK20 expression was still observed in these controls suggesting no technical issues. Assessing expression with immunofluorescence on the transverse plane may be more optimal for CK20. The absence of CK20 in the urothelium component may be due to co-culture effects altering expression in adjacent cells, incomplete differentiation of umbrella cells in this set of experiments, or shedding of the top layer during the co-culture or the ‘IHC sample preparation process.

Assessing cadherin expression is crucial as members of this family play a role in invasion and metastasis (34). E-Cadherin is known to be expressed throughout the bladder urothelium at the tight junctions and is a natural tumour suppressor (46). Cancers with a decreased expression of E-Cadherin exhibit a more invasive and metastatic behaviour, with the abnormal upregulation of N-Cadherin. This is consistent with the epithelial-mesenchymal transition (EMT) process whereby epithelial cancers transform to a migratory mesenchymal- like phenotype (47). In our experiments, RT112 spheroids incorporated into 3D-UHU did exhibit some migratory cells, but strongly expressed E-Cadherin, consistent with their classification as a low-grade cancer model. Conversely, T24 spheroids incorporated into 3D- UHU displayed a more frequent migratory phenotype and had an increased expression of N- Cadherin, consistent with their high-grade cancer classification. Given that low-grade cancers are typically well-differentiated and express E-Cadherin, which strengthens cell-cell adhesion, it is likely that they can tolerate the bladder’s luminal conditions without needing to invade more deeply into the nutrient-rich areas of the bladder wall.

In addition, MMP-2 and -9 are recognised as important biomarkers for bladder cancer prognosis (48). Kudelski *et al*. looked at their expression in low- and high-grade cancer tissue and also in healthy marginal tissue taken from the same patient. They found that MMPs were present in all tissue types regardless of grade and disease status, but MMP-9 expression was enhanced compared with MMP-2 particularly in higher-grade tissues (49). In both low- and high-grade 3D-UHU-TU models, only the presence of both these MMPs were investigated, and enzymatic activity was not measured. Our models also demonstrated a variety of migration types, consistent with the variety known to occur (50, 51) and demonstrating the dynamic and versatile nature of 3D-UHU-TU. It should also be noted that we did not observe an invasion phenotype in our higher-grade model, but it is possible that more time is required for this to occur, or that our model is lacking some component that might facilitate this. Enhancing the model to incorporate deeper layers of the bladder (e.g. lamina propria and or muscle microtissue) alongside more time in co-culture might allow us to model invasion in the future.

As the 3D-UHU-TU model is grown and maintained in a 100% urine environment, it was important to test the effect of the urine on cancer spheroid proliferation. The Ki-67 data from the low-grade RT112 spheroids within 3D-UHU-TU matched their proliferative nature seen in growth curves measured from individual spheroids growing in microcavity plates. In contrast, Ki-67 expression of the high-grade T24 spheroids was low within the main body of the spheroid, yet high in migratory T24 cells that had actively detached from the spheroid. Given that Ki-67 has been implicated in bladder cancer metastasis (52), this result is consistent with that role. In the healthy 3D-UHU component, proliferation was also seen in the basal layers and some intermediate cells. This was expected, as the transitional urothelium regenerates from the basal membrane upwards, with progenitor basal cells responsible for replacing cells that have matured. In addition, rapidly proliferating intermediate cells play a crucial role in establishing the multiple layers of the urothelium (33).

To assess the efficacy of 3D-UHU-TU as a testbed for new treatments, 3D-UHU-TU models were treated with the commonly used chemotherapeutic MMC. MMC induces apoptosis by alkylating DNA and forming crosslinks to prevent DNA synthesis (53). DNA fragmentation was seen in all treated RT112 spheroids which suggests that MMC did induce apoptosis. Normally, morphological signatures of apoptosis are DNA fragmentation and membrane blebbing (54) and in the absence of ‘scavenger cells’, secondary necrosis occurs (55). As the 3D-UHU-TU models do not incorporate immune cells, there are no immune cells present to phagocytose the apoptotic cells; therefore secondary necrosis is more likely to occur in the models. In our experiments, intracellular contents seemed to be lysing out of both cancer spheroids, suggesting secondary necrosis.

In conclusion, the 3D-UHU-TU model represents a new generation of *in vitro* urothelial bladder cancer model including a low and high-grade cancer component embedded in a healthy micro-environment, which shows promise as a testing platform for therapies. Like all models, 3D-UHU-TU does have some inherent limitations, for example the lack of a blood supply, deeper tissue layers or an immune system. On the other hand, in recent years, great strides have been made in incorporating more complexity into micro-physiological systems (reviewed in (56)), suggesting that the 3D-UHU-TU prototype could be used a foundation for developing more advanced iterations. One exciting possibility, which we are actively pursuing, is the incorporation of patient-derived tumour spheroids in place of cell lines, potentially allowing a personalised medicine approach to drug screening. Also, as *in vitro* cell-based model technology becomes more evolved, 3D-UHU-TU with such patient material could serve as a tool to help further understand the disease at the molecular level.

## Acknowledgements

We thank the Engineering and Physical Sciences Research Council, the UCL Therapeutic Innovation Networks and the Rosetrees Trust for funding the project. We would like to acknowledge the Research Histology Services and the Histopathology Department at Great Ormond Street Hospital (GOSH) for aiding us with the H&E and immunohistochemistry work, which was supported by the National Institute for Health and Care Research (NIHR) GOSH Biomedical Research Centre. The views expressed are those of the author(s) and not necessarily those of the National Health Service, the NIHR or the Department of Health. We would also like to thank Nenna Kanu and Mark Linch for the kind donation of RT112 cells.

---

**Figure S1:**
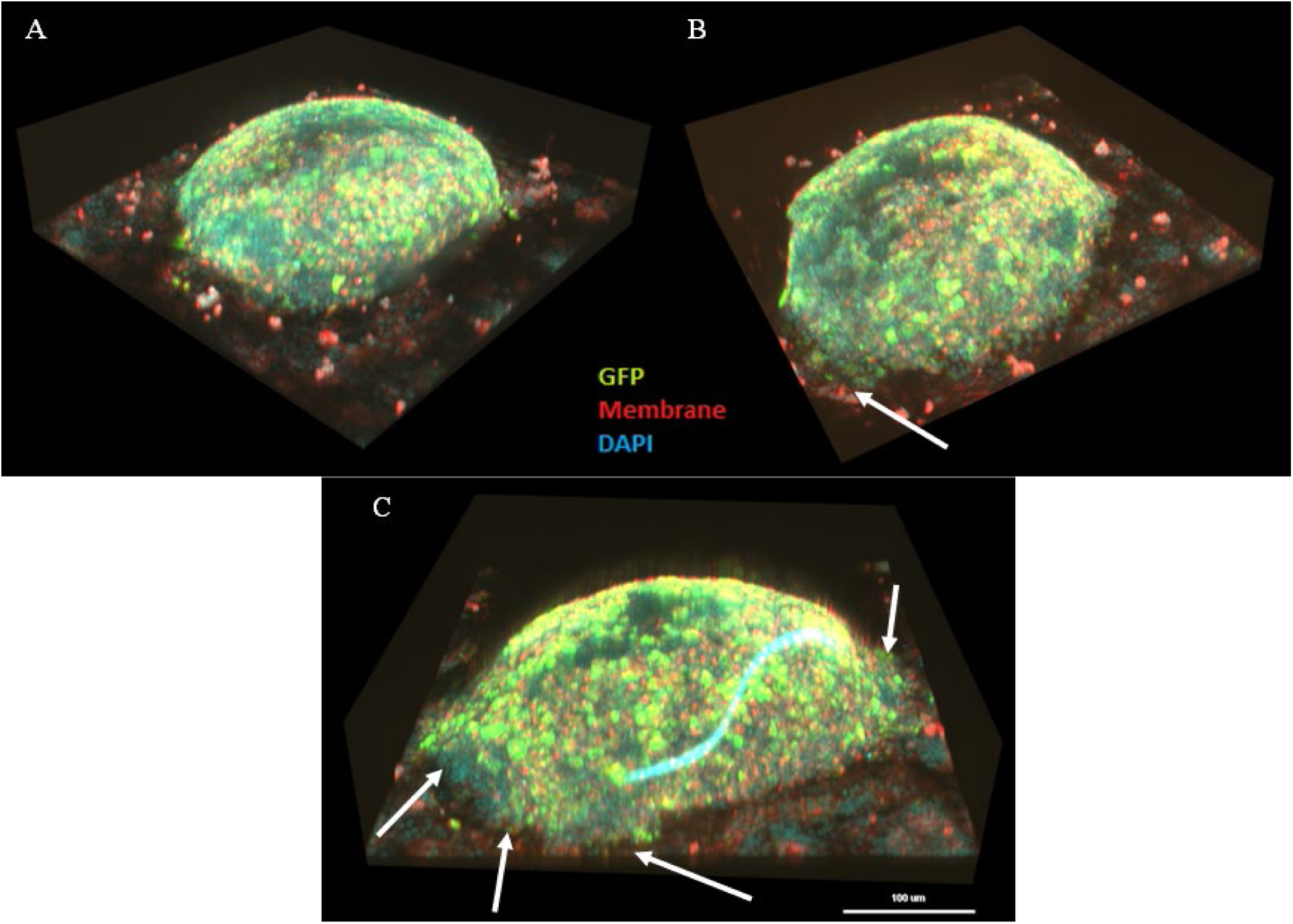
Light Sheet micrographs of RT112-GFP spheroids further boosted with GFP antibody staining (green) incorporated onto 3D-UHU (A) after 24 hours in 20% FBS; (B) after 24 hours in 20% FBS with intercalation of cancer cells (white arrows); (C) after 24 hours in 10% FBS with intercalation of cancer cells (white arrows). Blue strand is an unknown artefact. Membrane is stained with WGA-555 (red). Scale bar = 100 µm

**Table S1:** List of antibody details.

| Target/Stain | Host | Company | Product code | Concentration/<br>Dilution used | Other details |
| --- | --- | --- | --- | --- | --- |
| **Primary Antibodies** |  |  |  |  |  |
| **E-Cadherin** | Mouse | Invitrogen | 13-1700 | 10µg/ml | Monoclonal |
| **N-Cadherin** | Rabbit | Invitrogen | PA5-29570 | 1µg/ml | Polyclonal |
| **Ki-67 (IF)** | Mouse | Invitrogen | 50-5699-82 | 1µg/ml | Monoclonal;<br>Anti-human<br>eFluor™ 660<br>conjugate |
| **Ki-67 (Histo)** | Mouse | Leica | PA0230 | N/A | Bond RTU;<br>Monoclonal |
| **MMP-2** | Mouse | Invitrogen | 436000 | 2µg/ml | Monoclonal |
| **MMP-9** | Mouse | Invitrogen | MA5-15886 | 1:200 | Monoclonal |
| **CK7** | Mouse | Leica | PA0942 | N/A | Bond RTU;<br>Monoclonal |
| **CK20** | Mouse | Leica | PA0022 | N/A | Bond RTU;<br>Monoclonal |
| **GATA3** | Mouse | Sigma | 390M-14 | 1:300 | Monoclonal |
| **Phalloidin-555** | - | Invitrogen | A30106 | 0.04µM/ml<br>(1:500) | - |
| **Phalloidin-647** | - | Invitrogen | A30107 | 0.04µM/ml<br>(1:500) | - |
| **DAPI** | - | Invitrogen | D1306 | 1µg/ml | - |
| **Secondary Antibodies** |  |  |  |  |  |
| **Anti-Mouse 488** | Goat | Invitrogen | A11001 | 10µg/ml | - |
| **Anti-Rabbit 488** | Goat | Invitrogen | A11034 | 10µg/ml | - |
| **Anti-Mouse 647** | Goat | Abcam | ab150115 | 10µg/ml | - |
| **Anti-Rabbit 647** | Goat | Abcam | Ab150079 | 10µg/ml | - |

